# How Do Co-folding Models Organize Structural Information?

**DOI:** 10.64898/2026.09.06.749739

**Authors:** Minha Park, Shinwoo Kim, Seokhyun Moon, Hyeongwoo Kim, Gyuho Jeon, Woo Youn Kim

## Abstract

Co-folding models accurately predict biomolecular complexes, but how their internal representations support joint structure prediction remains unclear. We analyze the Boltz-1 trunk by decomposing its pair representation into intra-chain and inter-chain blocks and applying ablation, layer-wise representation geometry, and activation patching. Our results show that structural information is distributed across three streams: the single representation carries token-internal atomic detail, the intra-chain pair representation encodes chain-level geometry, and the inter-chain pair representation controls relative molecular arrangement. Across the trunk depth, representations follow a Mix–Compress–Refine trajectory, while linear probes reveal that intra-chain distance content is largely supplied by MSA preconditioning and inter-chain distance content is constructed progressively across Pairformer layers at each recycle. Apo–holo patching further shows that ligand-induced backbone changes are not fully resolved before diffusion; instead, the diffusion module reconciles partially inconsistent intra- and inter-chain constraints. These findings provide a mechanistic view of how co-folding trunks organize structural information and suggest more partition-aware designs for future biomolecular structure prediction models.

## 1. Introduction

Predicting biomolecular complex geometry is a central computational question in structure-based drug discovery. While individual protein folds provide a mechanistic basis for understanding biological activity, many drug-discovery questions concern how molecular partners interact. For those applications, the relevant prediction task extends from folding individual components to resolving the contacts and interfaces that organize them into a biologically relevant complex. Building on AlphaFold2’s success in single-chain protein structure prediction (Jumper et al., 2021), the AlphaFold3 family of *co-folding models* (Abramson et al., 2024) now predicts full biomolecular assemblies end-to-end from sequence, with a comparable open-source ecosystem rapidly following (Wohlwend et al., 2024; Passaro et al., 2025; ByteDance AML AI4Science Team et al., 2025).

Recent works have begun to investigate the internal workings of protein structure prediction models. For single-chain folding, both the model’s representations (Lu et al., 2026; 2025) and its components themselves (Ahdritz et al., 2024; Wang et al., 2025) have been examined. For co-folding, comparable analyses are far less developed (Ouyang-Zhang et al., 2025; Zhou et al., 2025; Migliorini et al., 2025), and the setting itself is more challenging: the Pairformer trunk additionally carries inter-chain pair blocks with no analogue in single-chain folding, so single-chain analysis methods do not transfer directly. As such, neither *what* these multichain-specific representations encode nor *how* they are built across layers has been substantively addressed.

We thus investigate these topics through ablation of the trunk’s representations, analysis of layer-wise representation geometry, linear probing of pairwise distances, and representation patching across apo/holo conformer pairs. From experiments designed around these methods, we make the following contributions:

- We show that the three types of representation encode distinct aspects of information: local atomic details, chain-level geometry, and relative orientation.
- We show how pairwise distance information evolves across the trunk and that representations follow a Mix-Compress-Refine-like depth profile (Skean et al., 2025).
- We show that the trunk’s representations are sometimes incomplete, with the diffusion module compensating.

## 2. Related works

### 2.1. Co-folding models

A *co-folding model* predicts the joint structure of a biomolecular complex from sequence alone, replacing the fold-then-dock pipeline with integrated inference over all chains (Jumper et al., 2021; Lin et al., 2023; Evans et al., 2021; Krishna et al., 2024). Modern co-folding models are highly inspired by AlphaFold3 (Abramson et al., 2024), and thus share the core network architectures: a *trunk* that constructs a per-token sequence representation and a pairwise representation; a *diffusion module* that decodes 3D coordinates from the trunk’s output. In a multi-chain assembly, the pair representation decomposes into intra-chain and inter-chain blocks, a structural distinction with no analogue in single-chain folding.

### 2.2. Interpreting folding models

Recently, single-chain folding models have been analyzed from multiple angles. Activation patching of ESMFold has shown that biochemical features such as charge are encoded as linear directions in the trunk’s representation, and that the trunk operates in stages: early biochemical signal propagation followed by late spatial feature formation (Lu et al., 2026). ESMFold’s input embedding has been shown to exhibit low intrinsic dimensionality (Lu et al., 2025). Training-dynamics analysis of AlphaFold2 retraining observed hierarchical learning of secondary before tertiary structure (Ahdritz et al., 2024), and Wang et al. (2025) showed that the trunk can be replaced by standard transformer blocks while retaining folding performance, suggesting that folding-relevant information can be encoded without domain-specific inductive bias.

For co-folding, PairSAE applied sparse autoencoders to the Boltz-2 pair representation and showed that it encodes token-level interpretable features that align with specific structural/biochemical concepts such as disulfide bonds and protease active sites (Migliorini et al., 2025). Architectural ablations also provide indirect insights into representational content: Pairmixer’s architecture omits in-trunk updates to the single representation while retaining strong performance, prompting a reinterpretation of the single representation’s informational role (Ouyang-Zhang et al., 2025); SeedFold finds that scaling the channel dimension of the pair representation is more effective than scaling trunk depth, highlighting the pair representation as a primary determinant of trunk capacity (Zhou et al., 2025). Feldman & Skolnick (2026) concurrently analyzed the co-folding model’s representations using several interpretability tools. While their work focused mainly on the role of MSA modules inside trunk in monomeric systems, ours investigates representations trunk underlying multimer modeling.

## 3. Methods

We analyze a co-folding trunk through four lenses organized along two axes, built on a chain-aware factorization of the pair representation into intra- and inter-chain blocks (**Z**_intra_, **Z**_inter_) and a frozen diffusion module shared across all interventions. Section 3.1 establishes the architecture and notation. The frozen diffusion module makes interventional lenses comparable: ablation (Section 3.2) and activation patching (Section 3.4) differ in whether they remove or substitute information, but both decode the modified representation through the same module. Observational lenses (Section 3.3)—representation geometry and linear probing— instead read content from a single unmodified forward pass.

### 3.1 Preliminaries: Co-folding architecture and notation

A modern co-folding model consists of two stages: representation generation and structure prediction (Abramson et al., 2024; Wohlwend et al., 2024; Passaro et al., 2025). In the representation generation, the major part is the *trunk*, which comprises a template, multiple sequence alignment (MSA), and Pairformer module. The first two modules encode structural and evolutionary priors into a token-level single representation and inter-token-level pair representation. Then, a Pairformer module iteratively refines both representations. Once the final representations are generated, a diffusion module decodes coordinates: it uses the single representation as per-token conditioning and the pairwise representation as an additive pair bias in the coordinate-denoising transformer (Abramson et al., 2024).

#### Notation

We write 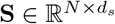 for the trunk’s single representation stack and 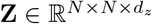 for its pair representation over *N* tokens; *s*_*i*_ and *z*_*ij*_ denote the corresponding per-token and per-pair slices. When the token set partitions into biomolecules, we decompose the pair array into a block-diagonal intra-chain subarray **Z**_intra_, pairs whose endpoints lie in the same chain, and off-diagonal inter-chain subarrays 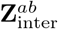, pairs whose endpoints lie in chains *a* and *b*. We write **Z**_inter_ when the chain pair is unspecified. Superscripts indicate the trunk depth at which an activation is read: **S**^(*l*)^ and **Z**^(*l*)^ are the *l*-th block output, and (**S**^(*K*)^, **Z**^(*K*)^) with *K* = 48 is the final state passed to the diffusion module.

#### The Pairformer trunk

Each block transforms 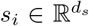 and 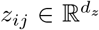 through four operations: triangle attention, triangle multiplication, a transition MLP, and a biased attention on the single representation. The first three operate identically on every (*i, j*) pair and hence on every block of the pair array regardless of chain membership. Triangle multiplication and triangle attention both update *z*_*ij*_ under a soft triangle-inequality constraint, propagating distance information across non-adjacent pairs. The transition MLP refines each representation position-wise, and the biased attention injects pair information into **S** by projecting **Z** into a bias term. More details are provided in Appendix A.1.

### 3.2. Ablation

Ablation is an interventional probe: we replace activations on a single forward pass and read out the effect of replacement under the same diffusion module. Interventions are applied independently to **S, Z**_intra_, and **Z**_inter_; this factorization localizes where geometric information is stored.

Given the final representations of the trunk (used as inputs to the diffusion module), we apply the following operations at the channel level. In other words, the operations are directly applied to the representation vectors *s*_*i*_ and *z*_*ij*_. The *zeroing* operation sets the chosen region to zero, collapsing both its content and its magnitude. The *mean* operation collapses the representations into the per-channel average *µ*_*c*_ so the channel retains its aggregate scale but loses position-specific structure. The *Gaussian* operation additionally computes the variance 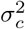, and samples each scalar element of the vectors from the Gaussian distribution 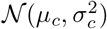. We refer to the mean-only and Gaussian forms collectively as *averaging ablation*, isolating structured-content removal from the scale effect of zeroing.

This follows the intervention logic common in mechanistic interpretability: one can perturb an internal representation during the model’s own forward pass and measure the resulting downstream behavior (Lu et al., 2026; Sun et al., 2024). In the co-folding setting, this can be assessed by the structure quality of the diffusion model outputs.

### 3.3 Representation geometry and linear probing

#### Representation geometry

Singular-value statistics of the activation matrix quantify how each layer compresses or expands its feature space (Queipo-de Llano et al., 2025; Skean et al., 2025). Each block’s pair representation is stacked into a real matrix *X*^(*l*)^ by reshaping 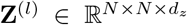 to 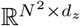, treating each (*i, j*) pair as a row and each channel as a column; for the single representation we use 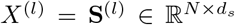 directly. Let *σ*_1_ ≥ *σ*_2_ ≥ · · · be its singular values and 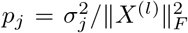 the normalized squared spectrum. We track three statistics that jointly characterize spectral concentration: the matrix-based entropy

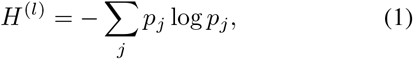

the stable srank 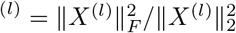, and the top-1 singular-value mass 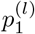. We compute these statistics at every layer in the trunk—the input embedder, the MSA module, and all 48 Pairformer blocks—and at every recycling iteration *r* ∈ *{*0, 1, 2, 3*}*, separately for **S, Z**_intra_, and **Z**_inter_.

#### Linear probing of pairwise distance

Linear probing fits a low-capacity decoder from hidden activations to a task target and quantifies the extent to which that target is linearly recoverable at each depth (Alain & Bengio, 2017; Belinkov, 2022). We fit a linear probe 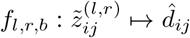 at every (layer *l*, recycle iteration *r* ∈ {0, 1, 2, 3}, pair-block partition *b {***Z**_intra_, **Z**_inter_}) configuration, where *l* spans the input embedder, the MSA module, and all 48 Pair-former blocks. The input is the symmetrized per-pair slice 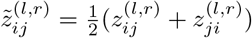 from the corresponding partition, and the target *d*_*ij*_ is the *C*_*β*_–*C*_*β*_ distance between protein residues *i* and *j* obtained from the top-1 (by confidence) of 25 diffusion samples per system; the probe therefore measures linear alignment between the pair representation and the geometry the diffusion module ultimately decodes.

Each probe is fit under 3-fold cross-validation; the details are given in Appendix A.3. We report the global pooled *R*^2^ as a function of (*l, r*), separately for **Z**_intra_ and **Z**_inter_, obtained by aggregating predicted–target pairs across all eval folds. The final-block *R*^2^ at the last recycle is the headline summary used to test whether final **Z** contains linearly accessible distance information.

### 3.4. Activation patching

Activation patching is a complementary interventional probe: rather than removing information from a single forward pass, we substitute activations from a donor forward pass into a recipient forward pass and read out the effect under the same fixed diffusion decoder. As in Section 3.2, patching is applied independently to **S, Z**_intra_, and **Z**_inter_ to localize the source of decoded geometry.

## 4. Results

We first briefly introduce our settings for each experiment in Section 4.1. To investigate the contribution of representations in structure prediction and their evolution through the trunk, we designed two targeted experiments: ablation (Section 4.2) and layer-wise analysis (Section 4.3). Building on the two preceding results, we designed a final experiment to assess where Boltz-1 stands relative to the ultimate goal of co-folding models: accurately predicting large conformational changes conditioned on the presence of a ligand. Concretely, we used activation patching to test whether the model’s internal representations encode the interaction between the two molecules (Section 4.4).

### 4.1. Experimental designs

#### Datasets

For the experiments, we used two datasets: the CASP set and ligand-induced conformational change set. The CASP set is a subset of public CASP15 and CASP16 data, selected to cover complexes that contain only protein and ligand molecules. CASP15 subset was filtered from the Boltz-1 evaluation set, and CASP16 subset was curated by ourselves. The ligand-induced conformational change set was specifically constructed by collecting paired systems with large protein conformational deviation for ligand-unbound (apo) and ligand-bound (holo) crystal structures. To this end, we implemented the ConfBench dataset proposed by NeuralPlexer3 (Qiao et al., 2026), as it was not publicly deposited. We chose to screen for apo-holo pairs where the target chain’s global RMSD lies between 1.5Å and 7.0Å ; the upper bound is imposed to restrict any implausible conformational changes. For more details about each dataset, see Appendix A.4.

#### Inference setup

All experiments used the Boltz-1 checkpoint distributed in the Boltz GitHub release v0.4.1 (Wohlwend et al., 2024), run on a single NVIDIA B200.

### 4.2. Where geometric content lives: ablation

We first analyzed the three representational components: **S, Z**_intra_, and **Z**_inter_. Recent architectural modifications to Boltz-1 remove updates to the sequence representation in the Pairformer module while retaining comparable end-to-end performance (Ouyang-Zhang et al., 2025). However, performance preservation under this architectural simplification does not imply that **S** is uninformative nor can it imply that **Z** is the sole carrier of information critical to structure prediction. This leaves open what information is encoded within **S** and **Z** which is used by the diffusion module. Furthermore, the setting in co-folding tasks that handles multimolecular systems calls for a further analysis into the **Z**_intra_ and **Z**_inter_ blocks and their contributions to structure prediction.

To localize the functional content of these representations, we used the ablation method explained in Section 3.2 and measured structure prediction metrics under each intervention. We used the CASP set defined in Section 4.1. For each target and ablation, we generated five diffusion samples and reported oracle metrics against the ground truth structures. Full metric definitions are provided in Appendix A.2; set-level statistics are shown in Figure 1, and additional results are deferred to Appendix B. We measured chain-LDDT, interface-LDDT, and LDDT-PLI to identify the effect of each ablation in forming chain-local geometry, inter-molecular arrangement, and protein–ligand contact quality.

**Figure 1.**
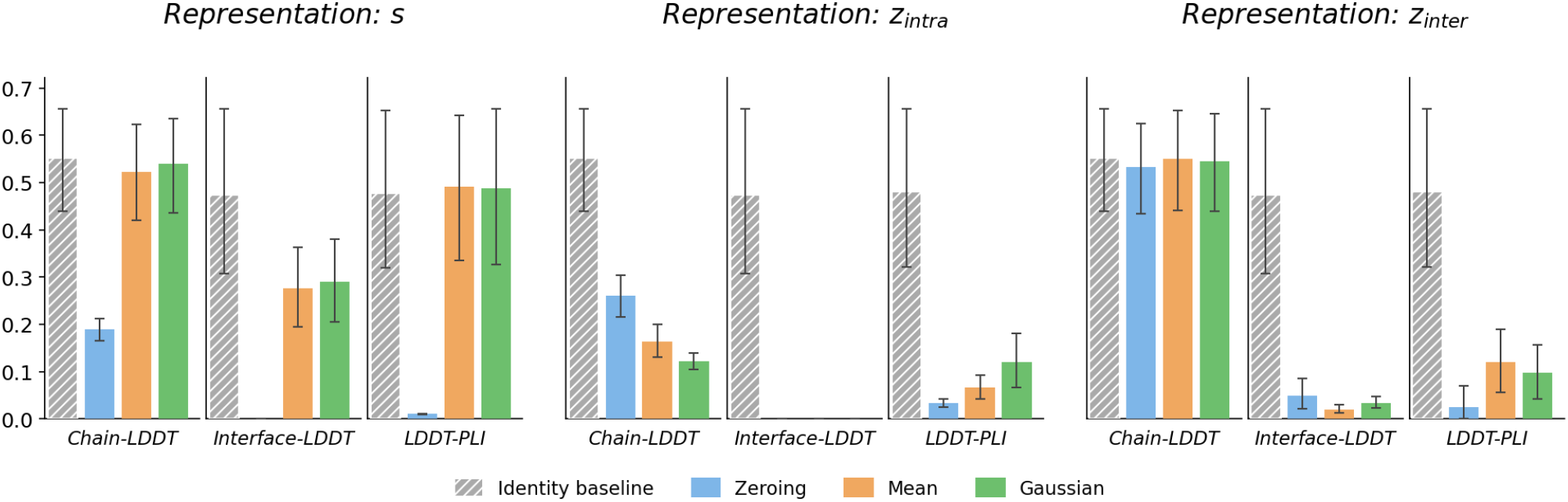
Set-level self-ablation outcomes on the combined CASP15+CASP16 multi-chain set. Bars show oracle summaries for zero, mean and Gaussian substitution of **S, Z**_intra_, and **Z**_inter_, with 95% bootstrap confidence intervals. The primary metric panels are chain-LDDT, interface-LDDT, and LDDT-PLI, separating chain-local fold quality, inter-chain arrangement, and protein–ligand contact preservation.

#### S contains token-internal atomic detail

Under mean and Gaussian ablations, both chain-LDDT and LDDT-PLI are maintained, whereas interface-LDDT deteriorates by nearly 40%. This indicates that once coarse single representation statistics are preserved, the unablated **Z** provides sufficient information to maintain chain-level geometry and most protein–ligand local scoring.

Inspecting the predicted structures further clarifies the role of the **S**. Under both mean and Gaussian ablations, the global chain remains largely aligned, but local atomistic geometry can become chemically invalid: Figure 2 shows that proline residues are incorrectly predicted as branched chains and not their true cyclic structures. Figures 5b and 5c also demonstrate poor side-chain structural prediction: although the relative placements of the atoms are correct, their interatomic distances are much smaller, causing the molecular visualization to render as a three-membered ring.

**Figure 2.**
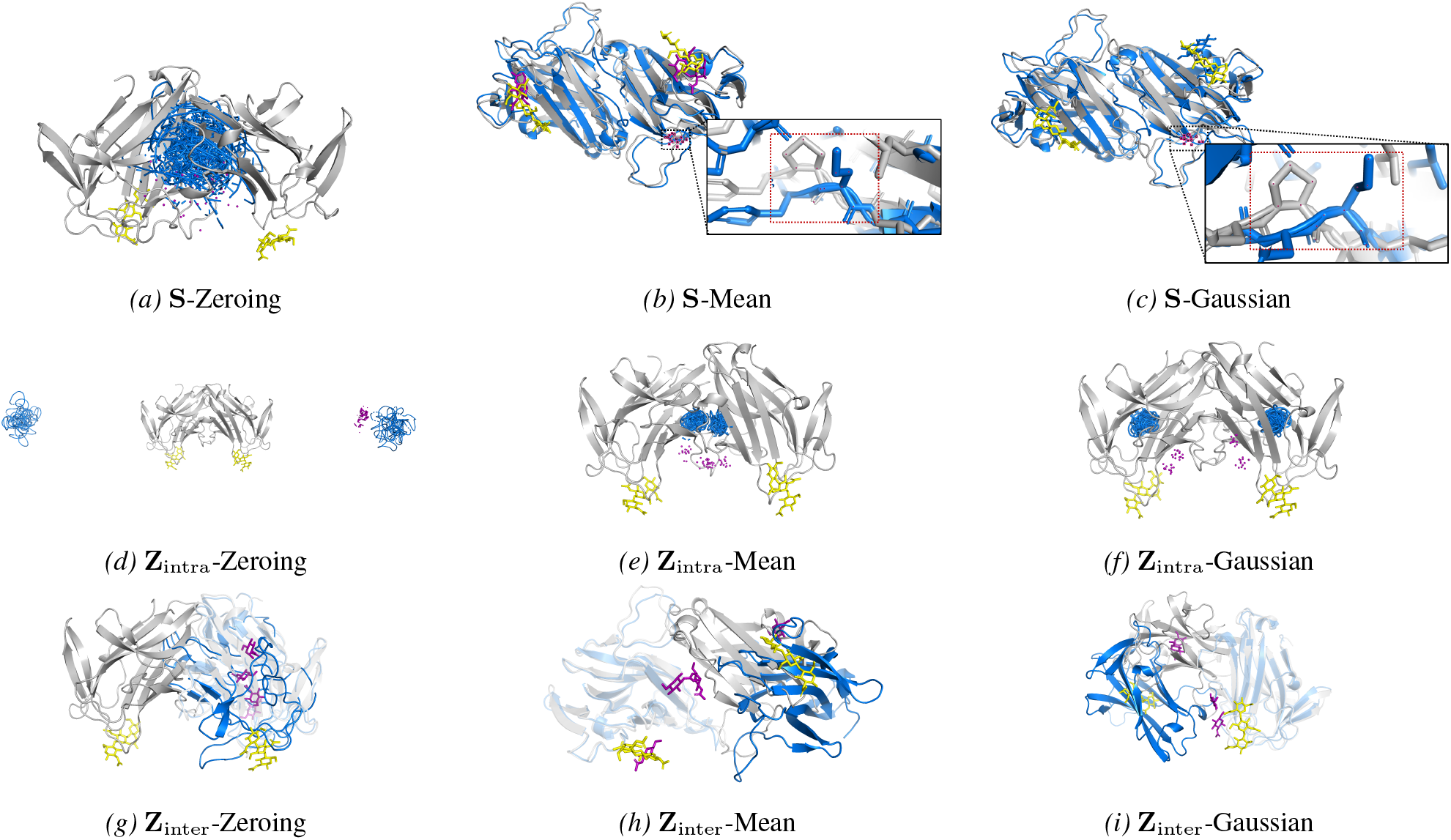
Case study on target T1187 from CASP15, which consists of 2 protein chains and 2 ligand molecules. The chains and ligands in the ground-truth structure are shown in gray and yellow, whereas those in the predicted structure are shown in blue and purple, respectively. In the **Z**_inter_ cases (g), (h), and (i), faded chains are aligned with each other.

Under Figure 1, zeroing **S** strongly damages all three primary metric families, but we note that this changes the activation scale and pushes **S** to an OOD-like region, rendering this ablation difficult to use as clean role-localization evidence. Such an effect of this ablation on the predicted structure is demonstrated in Figure 2.

**S** therefore acts as a conditioning stream that supplies the residue-tokens’ internal atomic detail rather than strictly defining chain-level geometry or arrangements.

#### Z_intra_ encodes chain-level geometry

Ablating **Z**_intra_ produces the strongest chain-local damage: under the averaging ablations, chain-LDDT drops by 70.2% and 77.9%. Mean and Gaussian substitution do *not* rescue chain folds, even though channel marginals (and, for the Gaussian variant, second moments) are preserved, as shown in Figure 2e and Figure 2f. Therefore, we conclude that **Z**_intra_ carries chain-level structural information.

#### Z_inter_ mediates relative positioning of molecules

Ablating **Z**_inter_ in any scope preserves chain-LDDT while degrading interface-LDDT and LDDT-PLI as shown in Figure 1. These results indicate that while the per-chain structures are preserved despite the ablation, the relative positioning of these molecules is broken. Figures 2g to 2i visualize this failure mode, and Figures 5g to 5i. This selective degradation of prediction performance indicates that **Z**_inter_ primarily encodes inter-chain structural information rather than intra-chain folding information.

### 4.3. How pair representations are built across trunk depth

The ablation results indicated that the final pair representation encodes geometric information. How is this content built up through the trunk at layer- and recycle-levels? We approached this from two complementary angles.

#### (i) How do Z_intra_ and Z_inter_ develop across the trunk?

The limit case is architecturally informative: if intra content were fixed by the MSA module and inter content added only later in the Pairformer, the trunk would amount to fold-then-dock under shared weights, a factorization that specialized single-chain and docking modules would express more directly. Locating where inter-chain content emerges therefore tests how cleanly this decomposition holds.

#### (ii) Does the layer-wise representation geometry follow the patterns in deep transformer stacks?

Representation geometry has recently been used to characterize layer-by-layer organization in large language models, revealing a characteristic MCR pattern across depth (Queipo-de Llano et al., 2025; Skean et al., 2025). These analyses, however, have been confined to standard self-attention. The Pairformer replaces it with triangle attention and triangle multiplicative updates across 48 blocks; whether the MCR pattern persists speaks to whether it is tied to attention form or is a generic property of information flow through self-attention mechanisms.

We address both questions with two observational analyses on the trunk’s layer outputs: representation geometry and linear probing based on the strategies defined in Section 3.3. We used a subset of the CASP set defined in Section 4.1 which includes only proteins.

#### Distance accessibility: Z_intra_ is MSA-saturated, Z_inter_ is built across Pairformer depth

Linear probes show distinct construction profiles for the intra- and inter-chain pair blocks (Figure 3a). As depicted in the final layer’s representation results in the linear probes, we first note that the final **Z** holds linearly accessible distance information. The **Z** that the diffusion module uses is linearly recoverable from **Z**^(*K*)^, strongly for 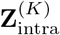 and more weakly for 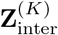.

**Figure 3.**
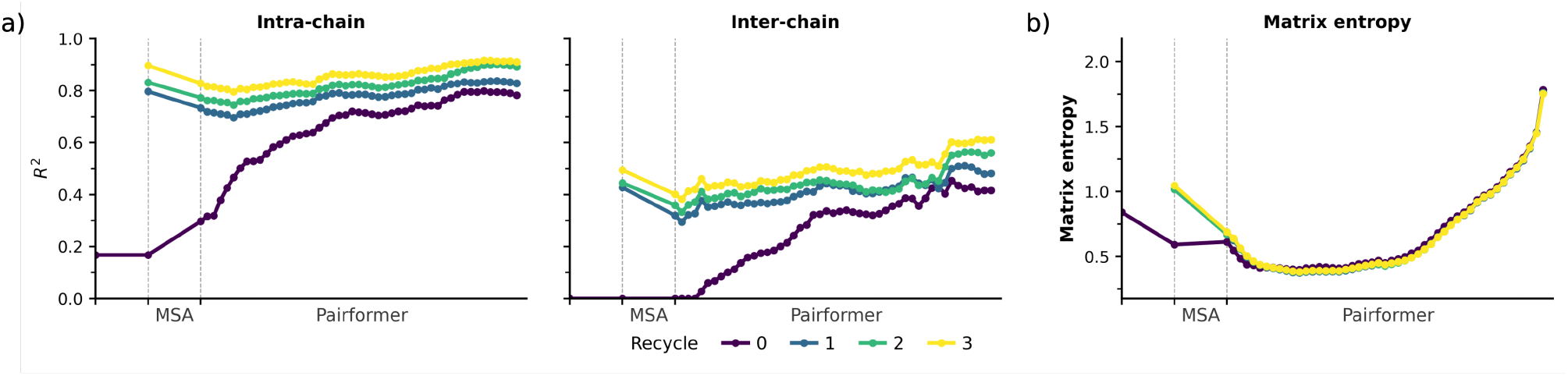
Layer-wise construction of pair representations across the trunk. *a)* linear probe global pooled *R*^2^ for decoded *C*_*β*_–*C*_*β*_ distance on **Z**_intra_ (middle) and **Z**_inter_ (right). The *x*-axis represents the pre-Pairformer region (z-init, post-recycle, post-MSA) and the 48 Pairformer blocks; recycles *r* = 0, 1, 2, 3 are colored separately. *b)* normalized matrix entropy of **Z** along the trunk.

We next examine how this distance content is built across trunk depth. In the first iteration (*r* = 0), the MSA module already supplied a substantial fraction of the intra-chain signal: **Z**_intra_ reached *R*^2^ ≈ 0.3 at the Pairformer entry, while **Z**_inter_ showed no correlation. Across the 48 Pairformer blocks, the *R*^2^ of both representations showed similar gains, so within a single forward pass the Pairformer updated intraand interchain pairs at similar rates. For *r* ≥ 1, **Z**_intra_ entered the Pairformer at *R*^2^ ≈ 0.80 and gained only marginally across the 48 blocks, whereas **Z**_inter_ improved more substantially yet still failed to close the gap to the intra ceiling. We interpret this divergence as asymmetric MSA preconditioning: under recycling, the MSA module transfers single-chain co-evolutionary signal predominantly into within-chain pairs, so that intra distance accumulates as an MSA-supplied asset that the Pairformer subsequently refines, whereas inter distance must be actively constructed across Pairformer depth in every cycle. The MSA-ablation collapse of pair representations in monomeric protein systems reported by Feldman & Skolnick (2026) for AlphaFold3 is consistent with this result.

#### MCR trajectory: the Pairformer compresses without losing distance content

Despite differing from a vanilla transformer encoder in using triangle attention and triangle multiplicative updates, the Pairformer trunk followed the Mix-Compress-Refine trajectory reported for language-model transformers (Queipo-de Llano et al., 2025; Skean et al., 2025): the MSA module mixed (entropy drops sharply), middle Pairformer blocks formed a compression valley, and late blocks re-expanded toward the diffusion handoff (Figure 3b). The same Mix-Compress-Refine phenomenon is visible in the stable rank and top singular value mass trajectories in Appendix B.3. The same valley emerged over what is effectively a single supervised target—pairwise distance content—in contrast with language-model multitask pretraining, suggesting that MCR is a property of self-attention information routing rather than of pretraining diversity. Distance accessibility did not collapse in layers where compression occured (Figure 3a): residual representational variance was removed while task-relevant content was preserved.

#### Recycling: two qualitatively different inputs share one set of weights

At *r* = 0 the input for the MSA module is low entropy with low distance accessibility (*R*^2^ ≈ 0 for **Z**_inter_), whereas at *r* ≥ 1 it has relatively high entropy and is near distance-saturated for **Z**_intra_ (Figure 3). Although the representations after each recycling step differ in information content, the same weights and normalizations are applied to them. Despite the fact that recycling actually improves the prediction performance, this entangles a build-up role at *r* = 0 with a refinement role at *r* ≥ 1 inside one module and forces the trunk to infer the regime implicitly from the input distribution, suggesting an architectural inefficiency that an explicit recycle-stage conditioning could relieve.

### 4.4. How does the trunk capture ligand-induced structural changes?

Many biomolecular complexes involve ligand-induced rearrangements of the protein backbone, so the chain-local geometry of the holo structure cannot always be inferred from the protein sequence alone. Whether a co-folding model handles this dependence is identifiable only where a ligand induces a large structural shift of the protein; in nearly rigid systems, a model that routes ligand-dependent information into the protein representation is indistinguishable from one that does not. We therefore conducted another analysis on protein–ligand systems with large backbone rearrangement between their apo and holo states using the ligand-induced conformational change set introduced in Section 4.1.

The previous two experiments indicate that co-folding trunks use **Z**_inter_ to encode binding-site and intermolecular contact geometry and **Z**_intra_ to express chain-local structure. Based on these observations, we hypothesized that protein representations should encode intra-chain distance information of the ligand-induced conformation in **Z**_intra_.

We tested this hypothesis with an apo/holo **Z**_intra_ patching protocol. We run independent inferences for the apo (ligand-unbound) and holo (ligand-bound) structures of the same protein sequence and obtain each trunk output’s (**S, Z**) pair. We then swaped the protein-block **Z**_intra_(*P, P* ) between the two trunk outputs in place. The h → a direction replaces the apo trunk’s **Z**_intra_ with the holo trunk’s, leaving the rest of the apo representation intact. The a → h direction is the mirror swap. Each patched representation is decoded through the diffusion module. We measured the protein chain’s backbone RMSD against the same system’s apo and holo predictions. The set of protein–ligand systems used for this analysis and per-system details are deferred to Appendix A.4.

#### The Pairformer does not fully separate information between Z_intra_ and Z_inter_ for the large conformational change set

Figure 4 reports, per system, the predicted protein-chain backbone RMSD to the apo prediction (x) and to the holo prediction (y), with lines connecting the apo, holo, and patched points of the same system. In both panels the patched point lies between the two references rather than collapsing onto the donor; the same-system line is pulled off the apo–holo axis. The two directions are individually informative. In the left panel of Figure 4 (h →a, apo recipient with holo **Z**_intra_), the patched point fails to reach the holo corner, indicating that holo **Z**_intra_ does not by itself carry enough chain-internal distance information to fix the backbone to the holo conformation. In the right panel of Figure 4 (a → h, holo recipient with apo **Z**_intra_), the patched point shifts away from the apo corner toward the holo prediction, indicating that the retained holo **Z**_inter_ supplies partner-dependent constraints that still influence the realized protein conformation. Overall, the generated chain conformation still depends on partner-dependent constraints carried by **Z**_inter_, indicating that induced fit is only partially resolved within the Pairformer before diffusion.

**Figure 4.**
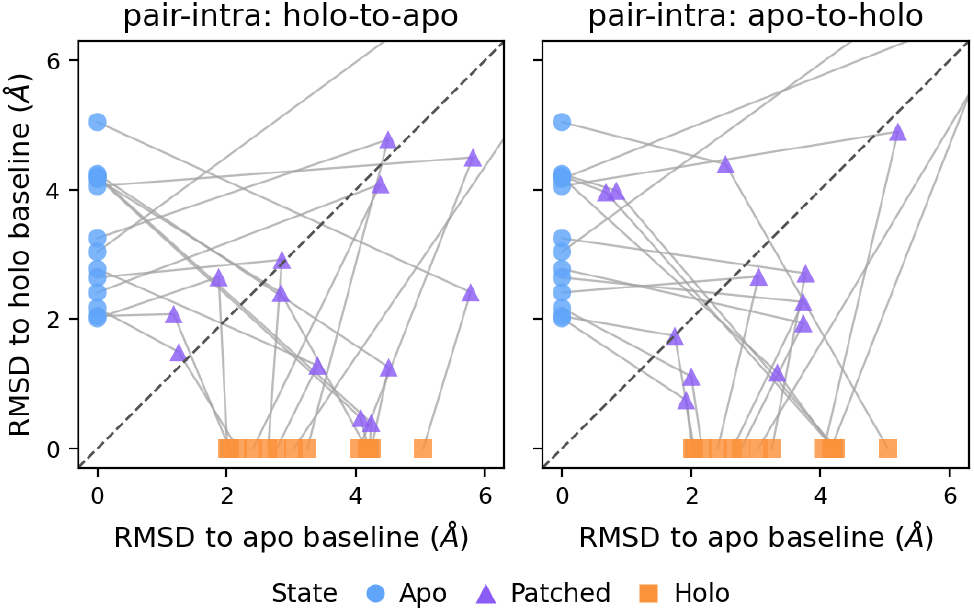
Activation patching results. Each panel plots, per system, the backbone RMSD of the predicted protein chain to the apo prediction (x) versus the holo prediction (y); lines connect the three points of the same system (apo, patched, holo). Left: holo-to-apo patching. Uses the **Z**_intra_ from the holo system and inserts into the corresponding apo pairwise representation. Right: apo-to-holo patching. Uses the **Z**_intra_ from the apo system and replaces the corresponding holo pairwise representation.

#### The diffusion module reconciles Z_intra_ and Z_inter_ rather than decoding a single distance field

The standard architectural decomposition in co-folding models treats the Pairformer as a pair-distance encoder and the diffusion module as a coordinate decoder, with the implicit assumption that the encoder produces a single internally consistent distance field for the decoder to realize. The patching outcome challenges this simple view: the Pairformer’s output is not internally consistent at the chain level, so the diffusion module reconciles partial and potentially inconsistent constraints from **Z**_intra_ and **Z**_inter_, together with the noisy coordinate state, rather than decoding one distance field.

## 5. Discussion

We combined three complementary views on the Boltz-1 trunk: ablation of the final representations (Section 3.2), layer-wise observational analysis of representation geometry and linear distance decodability (Section 3.3), and activation patching across apo/holo conformer pairs at the final representations (Section 3.4). Together they characterize *where* structural information is stored across **S, Z**_intra_, and **Z**_inter_, *how* that information is built across trunk depth and recycling, and furthermore *whether* the representation captures the ligand-induced large conformational changes in proteins. We summarize the main findings below and then discuss the model-design implications they suggest.

### 5.1. Main findings

- **Information is partitioned across the three representational components. S** carries token-internal atomic detail; **Z**_intra_ encodes chain-level geometry; **Z**_inter_ mediates inter-molecular arrangement.
- **The Pairformer trunk follows a Mix–Compress– Refine trajectory** despite using triangle attention and triangle multiplicative updates rather than standard self-attention, with task-relevant distance content preserved through the compression valley.
- **Recycling accumulates information, but inefficiently**. Linear probing shows little gain, yet the information gap between pre- and post-recycling stages remains substantial, motivating a more efficient design.
- **Ligand-induced structural change is only partially captured before diffusion. Z**_intra_ does not carry sufficient chain-internal information to fix the backbone to the bound conformation on its own, leaving the diffusion module to reconcile partial and possibly inconsistent constraints from **Z**_intra_ and **Z**_inter_.

### 5.2. Implications for co-folding model design

One direction concerns the entanglement between the build-up and refinement roles in recycling. Because the same Pairformer weights process inputs that differ qualitatively in entropy and distance saturation, the trunk likely under-utilizes its capacity at both stages. An explicit recycle-stage conditioning, implemented as a learned stage embedding injected into the residual stream or as gated stage-specific projections, would allow the network to allocate computation differently between initial construction and subsequent refinement without enlarging the parameter count substantially.

A second direction concerns the homogeneity with which the Pairformer treats **S, Z**_intra_, and **Z**_inter_ throughout the trunk. Our results indicate qualitatively different roles and information sources across the three: **Z**_intra_ is largely MSA-supplied and only marginally refined across Pairformer depth, **Z**_inter_ is actively constructed across depth at every recycle, and **S** acts as a conditioning stream of token-internal atomic detail rather than a vehicle for iteratively refined geometry. The current Pairformer applies identical triangle operations to all pair tokens, allocating equal capacity to a near-saturated intra block and a still-developing inter block. Decomposing the trunk’s pair operations along the intra/inter boundary, with lighter intra and heavier inter pathways, would align allocated compute with information growth, consistent with reports that pair-channel scaling outperforms depth scaling for trunk capacity (Zhou et al., 2025), and would also reduce the *O*(*N* ^3^) cost of triangle operations to a sum over smaller intra- and inter-chain blocks. Any such decomposition must preserve a mechanism for cross-partition information exchange, since our patching results show that intra geometry depends on partner-dependent constraints carried by **Z**_inter_.

## 6. Limitations

All experiments use Boltz-1; the chain-aware **Z**_intra_/**Z**_inter_ decomposition transfers directly to any Pairformer-style trunk, but reported magnitudes and depth-dependences remain model-specific until verified across families. Generalization beyond the integrated CASP15 and CASP16 multi-chain set to RNA, large multi-chain assemblies, and conformational-ensemble settings is not established here.

Ablation identifies what is destroyed by removing a representation, not what suffices to recover it; a representation written redundantly with **Z** would appear silent under **S** patches, so we do not claim **S** is uninformative in general. The layer-wise observational analyses do not share this destruction-asymmetry caveat: spectral and linear-decodability analyses speak to representation content directly. They introduce a complementary caveat: information presence does not entail information use. Averaging ablation numbers are conditional on the calibration-set protocol of Appendix A.

## Acknowledgements

This work was supported by the Basic Science Research Program through the National Research Foundation of Korea (NRF), funded by the Ministry of Science and ICT (Grant No. RS-2023-00257479). It was also supported by the Ministry of Science and ICT(MSIT), Republic of Korea, through the National IT Industry Promotion Agency(NIPA) (Grant No. PJT-26-100009).

## A. Method Details

This appendix collects the full equations, metric definitions, ablation details, and per-system dataset descriptions deferred from Section 3.

### A.1 Pairformer operations in full

Each Pairformer block consumes a single representation 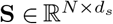 and a pair representation 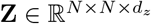 and emits an update of both streams (Jumper et al., 2021; Abramson et al., 2024; Wohlwend et al., 2024).

#### Triangle multiplicative update

The outgoing variant computes, for each pair (*i, j*),

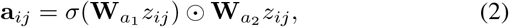

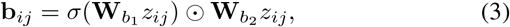

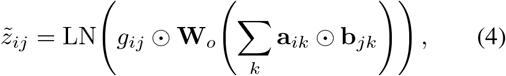

where *σ* is sigmoid, LN denotes layer normalization, *g*_*ij*_ = *σ*(**W**_*g*_*z*_*ij*_) is a sigmoid gate, and 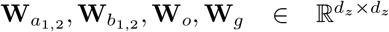 are learned projections. The incoming variant swaps the summation to ∑_*k*_ **a**_*ki*_ ⊙ **b**_*kj*_.

#### Triangle attention

The outgoing variant computes

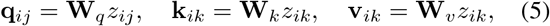

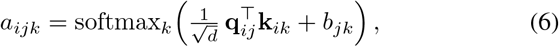

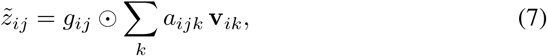

with learnable pair bias *b*_*jk*_ and sigmoid gate *g*_*ij*_. The incoming variant substitutes *kj* edges for *ik* edges and redirects the softmax accordingly.

#### Transition block

A position-wise two-layer MLP with expansion factor four and a gated activation (sigmoid-gated linear unit) is applied independently to every *z*_*ij*_ and every *s*_*i*_:

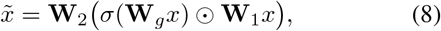

with *x* ∈ *{s*_*i*_, *z*_*ij*_*}* and residual connection.

#### Single-sequence update

After the pair operations, **Z** is read back into **S** via an outer-product mean followed by a multi-head self-attention layer biased by **Z**; details follow the AlphaFold-3 reference implementation (Abramson et al., 2024).

### A.2. Metric definitions

We collect here the structural-quality metrics used throughout Section 4. Unless stated otherwise all superpositions use OpenStructure’s chain-aware alignment, and all metrics are computed on full-atom predictions. The main self-ablation metrics are chain-LDDT, interface-LDDT, and LDDT-PLI.

- **Chain-LDDT**: the local distance difference test over all atom pairs within 15 Å, computed superposition-free as the fraction of inter-atomic distances preserved within distance tolerances *{*0.5, 1, 2, 4*}* Å and averaged over the four tolerances. We report the global mean over all atoms in the predicted complex.
- **Interface-LDDT**: the same LDDT computation restricted to inter-chain atom pairs whose ground-truth distance is within 15 Å . It is sensitive to chain arrangement but insensitive to chain-internal fold quality, which is what makes the chain-LDDT/interface-LDDT contrast useful for separating fold-level damage from arrangement-level damage in Section 4.2.
- **LDDT-PLI**: the variant of LDDT computed on protein– ligand atom pairs, used as the primary measure of ligand placement and protein–ligand contact preservation.

### A.3. Linear probing details

At each representation block *l* we fit a ridge regression probe 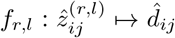 on the symmetrized channel features of *z*^(*l*)^. The ridge penalty is set by a scale-aware default *λ* = *α*_scale_ · mean(diag *S*_*xx*_) with *α*_scale_ = 10^−6^. For each fold, a probe is trained on 28 systems of the multi-chain set and evaluated on the remaining 14 systems. The target is the *C*_*β*_– *C*_*β*_ distance in the best-of-25 diffusion samples (ordered by confidence), not the crystal distance. The reported metrics are global pooled held-out *R*^2^, computed per block depth *l* and per pair-block partition 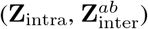.

### A.4. Details about the dataset

#### The CASP set

The CASP15 subset is taken from the evaluation set used in the Boltz-1 paper; both the YAMLs and the MSA files reported by Boltz is used. The CASP16 subset is a self-curated set where the experimental structures are released on the RCSB Protein Data Bank before March 11, 2026. The relevant MSAs were generated by the base AlphaFold 3 pipeline. After filtering problematic systems and removing targets that caused OOM errors, the set comprises 41 systems spanning multimolecular assemblies.

#### The ligand-induced conformational change set

The donor and recipient pairs used in Section 4.4 were selected to be a curated subset of the self-implemented ConfBench dataset (Qiao et al., 2026), which in turn is based on the PLINDER database (Durairaj et al., 2024). Of these apoholo pairs, we cluster the apo-chain sequences using MM-seqs2 and sample the apo-holo pairs from different clusters to ensure diversity in the systems predicted. Furthermore, we restrict to systems where the PDB-deposited bioassembly reports systems with less than 1,000 residues. Regardless, the structure prediction is on the individual chain and ligand of interest specified by PLINDER’s apo-holo pair annotations; each Boltz-input YAML for the apo system thus should contain a single protein sequence, and each YAML for the holo system should contain a single protein sequence and a corresponding ligand.

## B. Additional Results

### B.1. Self-ablation results on a protein–ligand system

This section provides additional qualitative evidence supporting the analysis in Section 4.2, where we examine where different types of geometric information are encoded through self-ablation. While Figure 2 focuses on a protein– protein system, Figure 5 shows a representative protein– ligand system.

The enlarged views in the Figure 5b and Figure 5c cases show that ablation of *s* leads to inaccurate atomic-scale reconstruction of the protein chain, although the overall chain-level arrangement is partially retained. In contrast, ablation of *z*_intra_ causes a substantial failure in the overall structure prediction, indicating that intra-chain pair representations are critical for preserving the internal geometry of the protein. The Figures 5g to 5i further illustrate the role of inter-chain or protein–ligand interaction information. When the faded chain A structures are aligned, the predicted ligand positions deviate substantially from the ground-truth ligand positions. This suggests that ablation of *z*_inter_ disrupts the model’s ability to accurately predict protein–ligand interactions.

**Figure 5.**
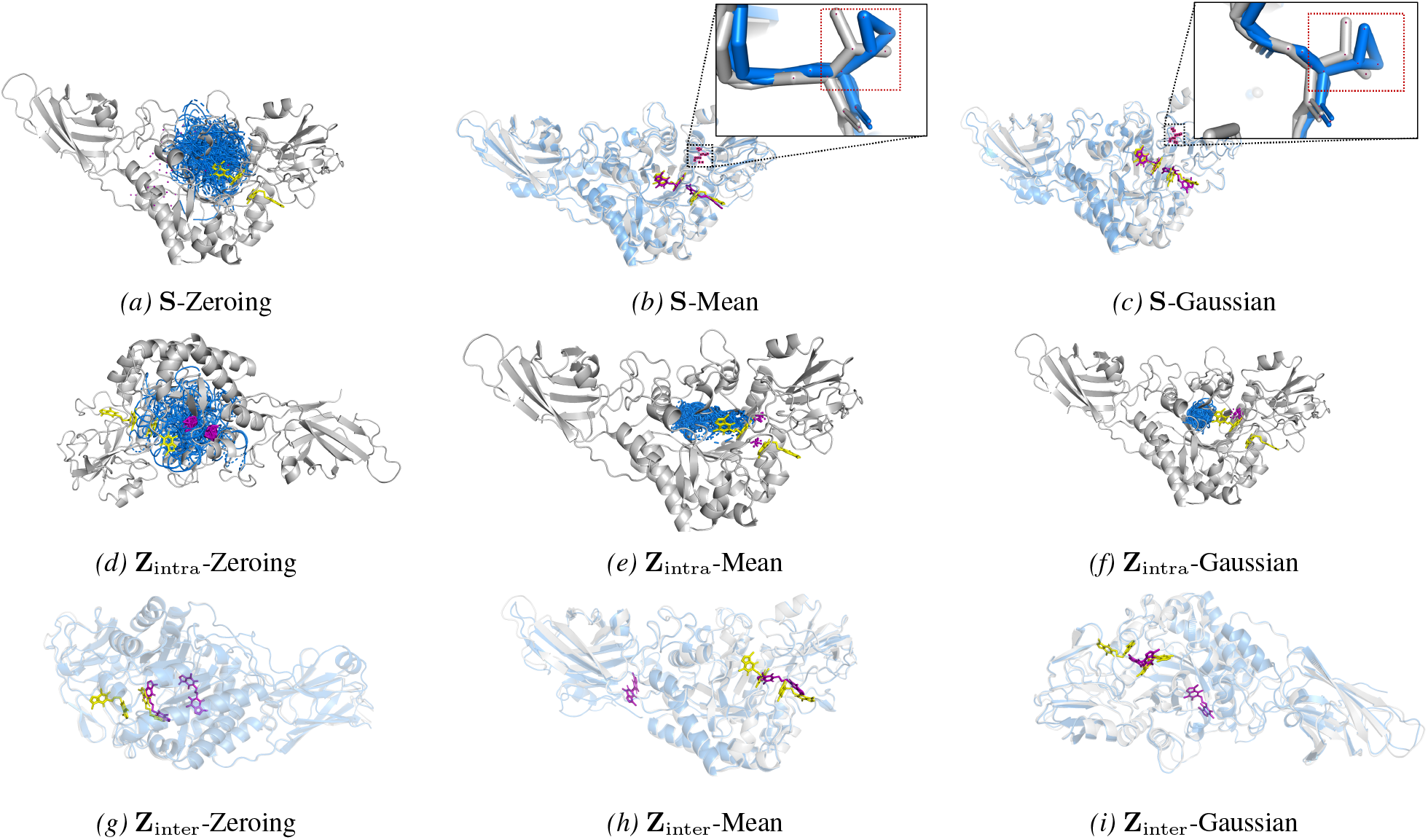
Case study on target T1188 from CASP15, which consists of 1 protein chain and 2 ligand molecules. The chains and ligands in the ground-truth structure are shown in gray and yellow, whereas those in the predicted structure are shown in blue and purple, respectively. In the Figures 5g to 5i cases, faded chains are aligned with each other.

### B.2. Top-1 self-ablation results

Figures 6 reports the corresponding top-1 measurements for the self-ablation settings shown in Figures 1. While the oracle results summarize the best prediction among sampled structures, the top-1 results are obtained from the model with the highest confidence score among the sampled predictions. Thus, the top-1 results provide a complementary view of the practical effect of each ablation setting under the model’s own ranking criterion.

**Figure 6.**
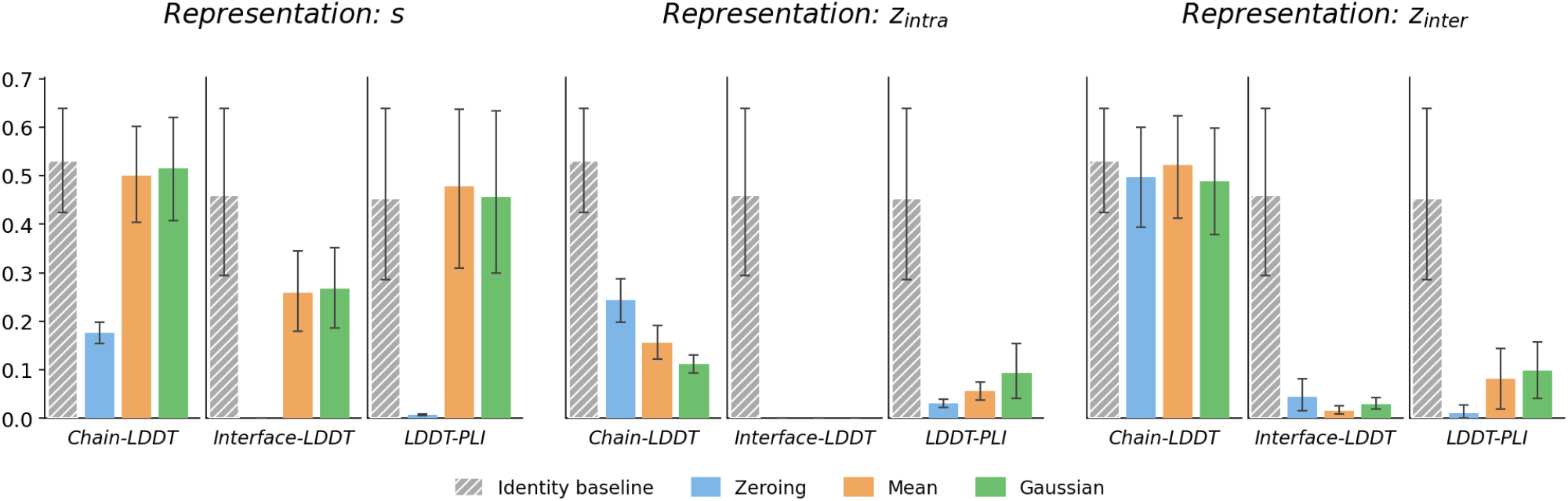
Set-level self-ablation outcomes on the combined CASP15+CASP16 multi-chain set of top-1. Bars show top-1 summaries for zero, mean and Gaussian substitution of **S, Z**_intra_, and **Z**_inter_, with 95% bootstrap confidence intervals. The primary metric panels are chain-LDDT, interface-LDDT, and LDDT-PLI, separating chain-local fold quality, inter-chain arrangement, and protein–ligand contact preservation.

### B.3. Additional spectral analysis of Z

Figure 7 extends the spectral analysis and shows Mix-Compress-Refine theory given by (Queipo-de Llano et al., 2025). We plot the stable rank and top singular value masses and show that they also follow the MCR story shown by the matrix entropy in the main paper (Section 4.3).

**Figure 7.**
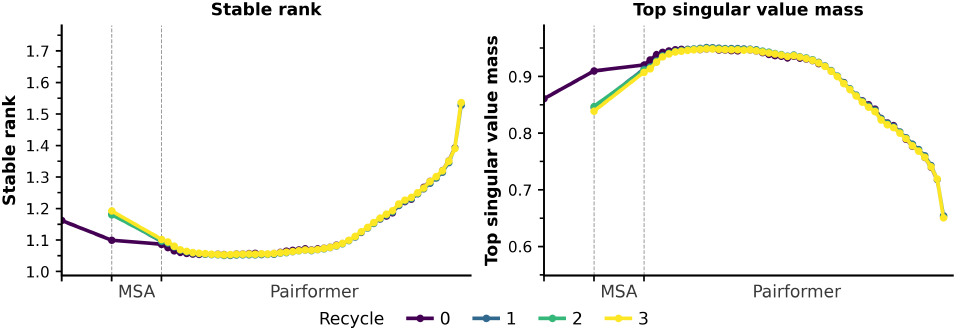
The flattened **Z** matrix’s stable rank and top singular value masses across the trunk.

## Notes

### Competing Interest Statement

The authors have declared no competing interest.

